# Repeated Activation of Gαq by neurotransmitters causes age-dependent accumulation of stress granules in *C.elegans*

**DOI:** 10.1101/2023.12.04.569992

**Authors:** Madison Rennie, Suzanne Scarlata

## Abstract

Gαq proteins mediate signals from neurotransmitters to transduce calcium signals. In PC12 cells, we have shown that Gαq stimulation results in retraction of neurites and induces the formation of stress granules that sequester two specific mRNAs, *Chbgb* and *ATP5f1b*. Here, we show that repeated activation of Gαq in *C. elegans* adversely reduces lifespan potentially through accumulation of stress granules and inefficient recovery from neurite retraction. In the absence of stimulation, we could not detect significant changes in number of stress granules from Day 1-15 worms. Single Gαq activation increases stress granule size through the enlargement of pre-formed particles with younger worms (Day 1-4) being more responsive than older (Day 8). Repeated Gαq stimulation impacts the number of particles through the assembly of nascent particles. Additionally, we a systematic rise in the number of AGL-1 stress granules with repeated Gαq stimulation suggesting that stress granules accumulate in neurons and sequester mRNAs. This idea is supported by immunofluorescence studies of ATP5f1b as well as changes in peristaltic speed. In addition to stress granule accumulation, we find that repeated Gαq activation results in age-dependent morphological defects in mechanosensory neurons. Taken together, studies show that repeated Gαq activation negatively influences the health of *C.elegans*.

## INTRODUCTION

Organisms are subject to a variety of environmental stresses throughout their lifetime, and the ability to respond to stress depends on age and fitness. *Caenorhabditis elegans* (*C. elegans*) are an attractive model system to study age-related responses to changes in environment due to their short life-span and ease of genetic manipulation (see (1) for review). Recent studies have screened for genes involved in *C.elegans* aging. These genes mediate protein homeostasis, energy production and mitochondria function as well as other cell functions (see (2-4)).

The short lifespan of *C. elegans* also makes them a good model system to study neurodegenerative diseases where the underlying biology and late onset in mammalian systems is difficult to study. In humans, many neurodegenerative diseases involve increased expression of proteins that form intracellular aggregates which interfere with neuronal function as seen in loss of cognition and movement (5), such as the transgene-encoded poly Q of Huntington’s disease or the α-synuclein of Parkinson’s disease (see (6)). The relationship between aging and protein aggregation may also be operative in *C. elegans* where it has been found that reducing the levels of aggregation-prone proteins increases the lifespan of worms (7, 8). Functionally, aging studies of worms show that sensory stimulation seems to remain intact with age, but muscular functions such as locomotion decline (9). While the genetic factors that contribute to C. *elegans* longevity have been uncovered (2), much less is known about their age-related responses to environmental stress on the cellular level.

Recent studies have found that muscarinic agonists, which stimulate the Gαq/ phospholipase Cb (PLCβ) signaling system, can reduce age-related decline of function (9). Our lab studies the Gαq/PLCβ pathway in cultured neuronal cells (e.g.(10)). This pathway is activated when neurotransmitters such as acetylcholine, serotonin, bradykinin, endothelin II and histamine bind to their specific G-protein receptor (11-13). This binding activates Gαq, which then activates PLCβ, setting off a series of events that result in increased intracellular calcium (12). We have previously found that in model cultural neuronal cells (PC12), Gαq activation causes irreversible retraction of neurites in the first 15 minutes of stimulation (14) and causes the cells to return to a stem-like state after 72 hrs post-stimulation (15). Alternately, in *C.elegans*, Gαq activation transiently ruptures neuronal connections and halts locomotion which then recovers after ∼30 minutes presumably due to endogenous growth factors (14).

Aside from this important lipid signaling function, we have found that Gαq activation impacts protein translation and stress granule formation in cultured cells (16-19). This activity is carried out through the cytosolic population of PLCβ. Under basal conditions, cytosolic PLCβ binds to proteins involved in RNA-induced silencing (16, 20) and stress granule formation (18). Activation of Gαq on the plasma membrane causes relocation of the cytosolic PLCβ population to the plasma membrane and the release of its binding partners. The release of stress granule proteins allows for their aggregation into stress granules (18).

Stress granules are halted translation complexes that form when cells are subjected to harsh and/or non-physiological environmental conditions such as nutrient deprivation, oxidative or toxic agents, and temperature variations (21-23). The formation of stress granules enables cells to shift protein synthesis to other proteins that allow the cell to better accommodate the stress. Typically, stress granules are found to be dynamic structures, where they form and dissipate within 1-2 hrs after a stress event (24). Aberrant stress granules are associated with aging (25). While stress granules form under adverse conditions, we also find that stress granules form when Gαq is activated. We have found that in PC12 cells, the stress granules that form as a result of Gαq activation sequester two specific mRNAs, a subunit of ATP synthase (*ATP5f1b*) and a protein that mediates vesicle formation (*Chgb*), in contrast to stress granules formed upon heat shock which sequester many different and non-specific RNAs (26). Because Gαq signaling is a normal physiological event in many species, it is possible that stress granules formed upon Gαq activation will accumulate if re-activation occurs before complete disassembly.

Here, we characterized the presence of stress granules in *C. elegans* neurons as a function of age, and with single and repeated Gαq stimulation. While aging does not significantly increase the number of stress granules, worm responses to Gαq varied with age, and importantly, we find that repeated Gαq stimulation results in the accumulation of stress granules and is associated with a diminished lifespan. Taken together, our studies show that repeated neurotransmitter stimulation may have detrimental impact on *C.elegans* health.

## RESULTS

### Repeated Gαq stimulation impacts *C. elegans* lifespan

Although the genes that transcribe Gαq and PLCβ are not associated with *C. elegans* longevity (see (27)), we have found that the gene that encodes a subunit of ATPsynthase, *ATP5f1b*, is sequestered in stress granules formed in response to Gαq activation. This sequestration may impact mitochondria health, which is closely associated with longevity (28).

We followed the lifespan of *C. elegans* after the Gαq/PLCβ pathway was activated once or multiple times. Worms were synchronized by bleaching (see Methods) and recordings began after all worms reached the L4 larval stage. Worms were either stimulated by carbachol for 30 min to activate Gαq or repeatedly (i.e. 30 min/day) throughout their lifetime. The results were compared to the lifespan of unstimulated worms. As shown in (**Fig. 1A)**, we find a detrimental effect of repeated Gαq/PLCβ activation on the worm lifetime after repeated stimulation as compared to control and single stimulation conditions.

**Figure 1.**
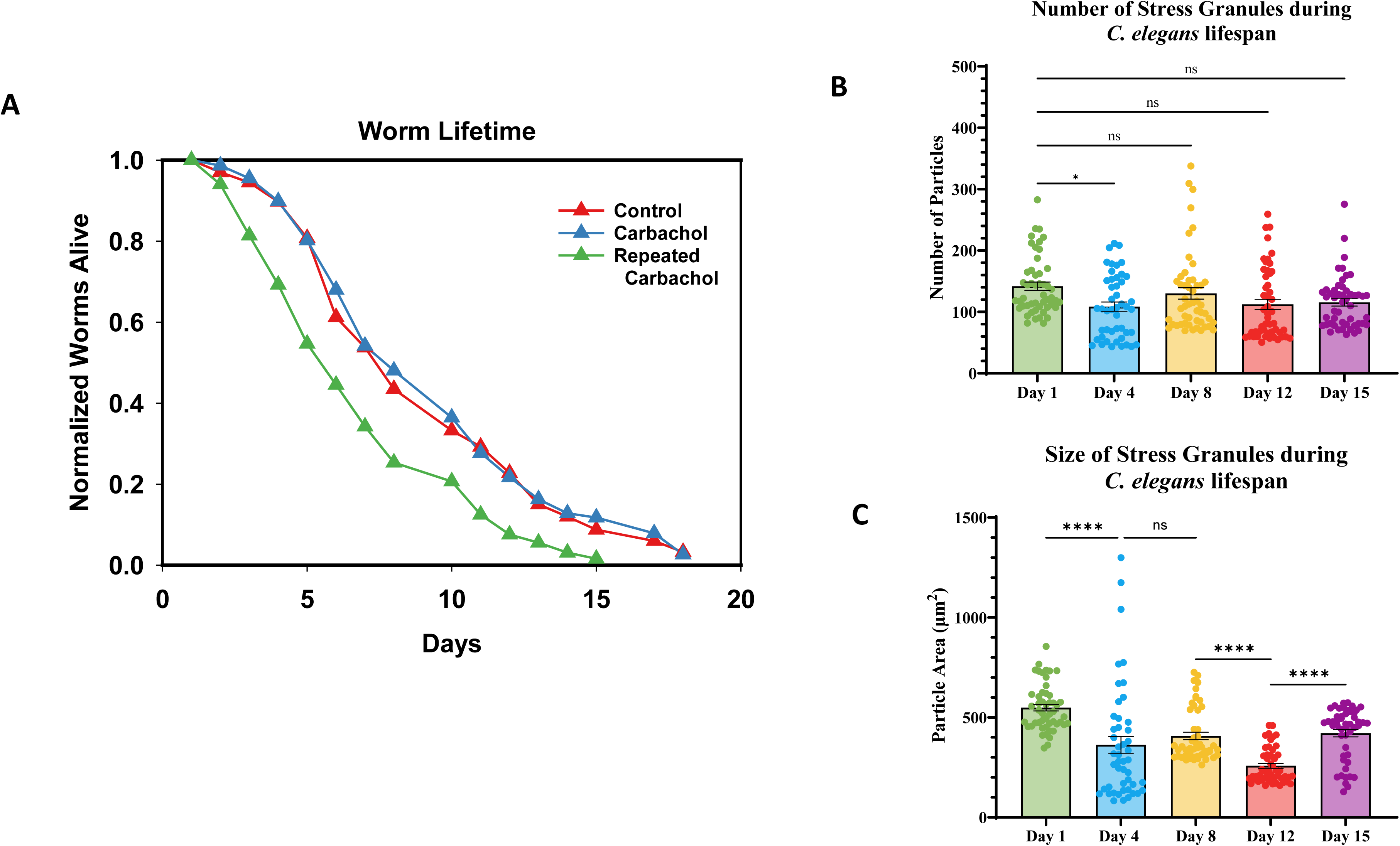
Survival and accumulation of stress granules of wild-type N2 C. elegans during their lifespan. **(A)** Survival results of worms under control and single and repeated Gαq stimulation where C. *elegans,* beginning at day 1. L4 larvae were exposed to 30min 1 mM carbachol at room temperature for 30 min, or repeated Gαq stimulation beginning at L4 larval stage until time of death). After carbachol exposure, worms were placed on NGM plates seeded with OP50 and were kept at 20°C. Alive worms were picked and moved to fresh plates daily, until time of death. For all conditions, n= 89-103 and N=3. Data was plotted using GraphPad Prism and analyzed using the Log-Rank (Mantel-Cox) test, where p= 0.0001. **(B-C)**. Age-dependent aggregation of G3BP1::gfp in the crossed strain JH3199::NM4397 held under basal conditions where C*. elegans* head neurons were analyzed at the first day of adulthood until day 15 of adulthood. Cumulative values of the number of observed particles and size of G3BP1::gfp particles were determined by confocal imaging under control conditions (see Methods).

### The number of *C. elegans* stress granules in mechanosensory neurons remain constant with age under normal growth conditions

Before assessing the impact of Gαq activation on stress granule formation, we first monitored how these particles change during the adult lifetime. We quantified the size and number of stress granules in confocal slices through organisms expressing eGFP-G3BP1, where dimerization of G3BP1 is a critical first step for the formation of stress granules (29). eGFP-G3BP1 was expressed in a crossed wormline that also expresses an RFP cytosolic marker for mechanosensory neurons (JH3199::NM4397). These worms allow us to view stress granules in the green excitation range (eGFP::G3BP1) by selecting neuronal regions in the red excitation range (mec-7p::mRFP) which label the mechanosensory neurons found in the head region of the worm. To analyze stress granules, we collected confocal images of the worm to create a series of optical slices spanning the width of the whole organism and identified particles ∼10µm^2^ or greater. The results are shown in **(Fig. 1B-C)**. We find that in adult worms, aged 1 to 15 days, the average number of stress granules in neurons remains constant, although the average size of the stress granules decreases with age, indicating aging is associated with reduced size of stress granules without a change in number.

### Assembly and disassembly of stress granules in *C.elegans* mechanosensory neurons with single and repeated Gαq stimulation

We followed the changes in size and number of G3BP1 stress granules after a 30-minute incubation with carbachol to stimulate Gαq, and 30-minute recovery after the worms were removed from the stimulus (**Fig. 2A-D**). Gαq stimulation increases both the size and number of stress granules in cultured cells (18) and we find increases in stress granule size in Day 1 worms (**Fig. 2A-B**). However, middle-aged and older worms show a reduction in both size and number suggesting a redistribution of preformed stress granules rather than the formation of new ones (see below). In cultured cells, stress granules recover in ∼30 minute but, in C. elegans, Day 1 worms stress granules continued to form, middle-aged worms showed increases in both size and number, and older worms showed a low number (**Fig. 2C-D**). These results show that the recovery process in worms is longer than in cultured cells.

**Figure 2.**
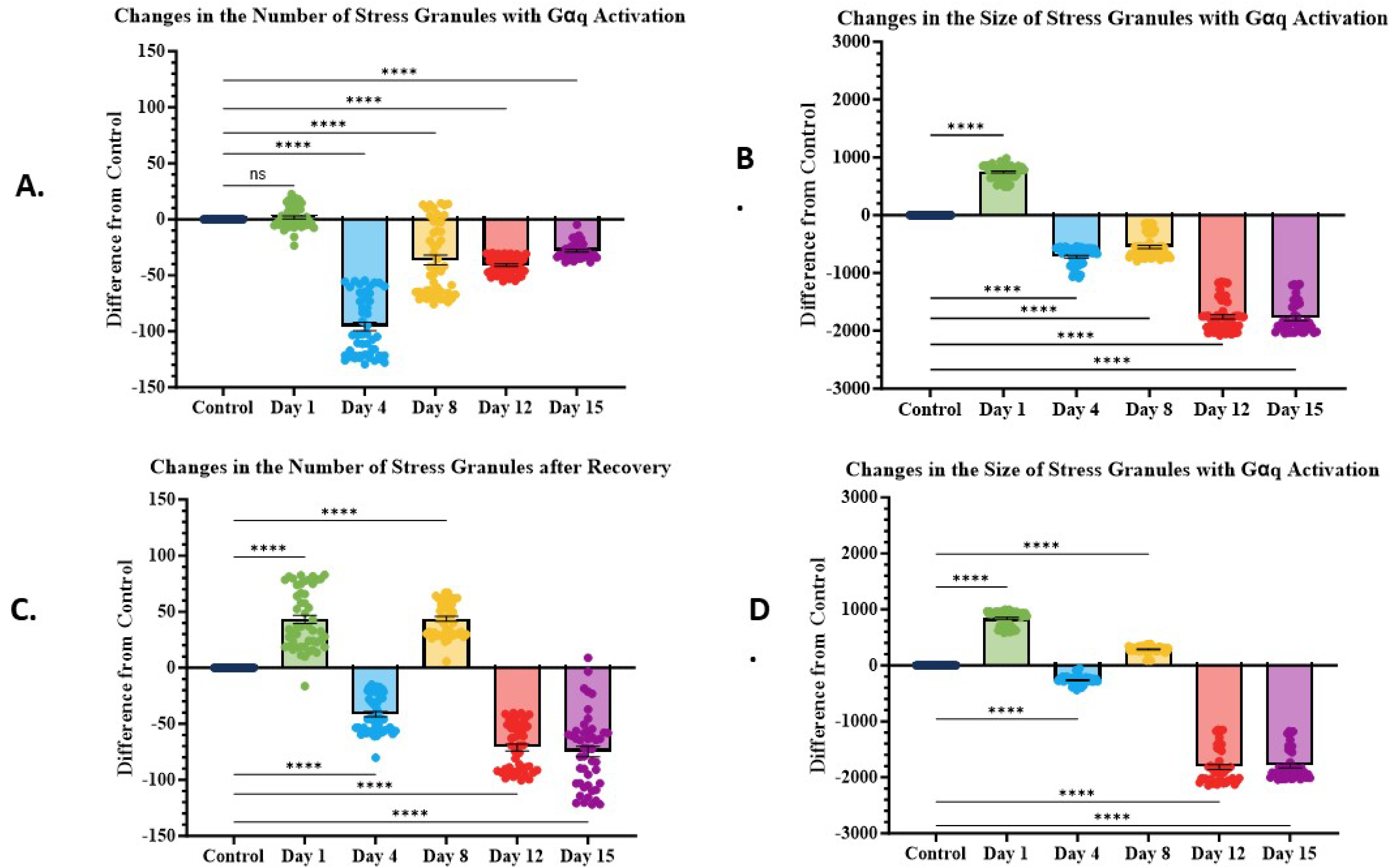
Age-dependent changes the assembly of G3BP1 stress granules in C. elegans neuronal cells upon Gaq stimulation and recovery. **(A-B)** Number and size of G3BP1 particles in worm head neurons (JH3199::NM4397) stimulated treated with 1mM carbachol for 30 min to stimulate Gαq, and **(C-D)** recovery in the number and size of the particles after 30 min on an NGM plate seeded with OP50. For all ages, n=10-15, where SEM and p-values are shown. Data was visualized using GraphPad Prism and analyzed using a One-Way Anova, where p-values are represented on the graph. (“ns” correlates to non-significant data points and **** correlates to a p-value < 0.0001.

### Monitoring the assembly and disassembly of Gαq- mediated stress granules

In cultured cells, activation of Gαq increases the size and number of stress granules through a mechanism involving the release of stress granule proteins bound to cytosolic PLCβ (18). In addition to following larger stress granules by confocal imaging of eGFP-G3BP1 in worm mechanosensory neurons, we monitored stress granule formation by following the onset of G3BP1 oligomerization using number and brightness (N&B) studies. N&B monitors the number of fluorophores in a diffusing species (30), and these values will increase with G3BP1 dimerization which initiates stress granule formation. In this analysis, the brightness and intensity values of the diffusing fluorescent particle are plotted and the points are enclosed in a selected area of the graph and compared to a non-aggregating control fluorophore, or control conditions. Increased aggregation is indicated if the points from the sample lie outside the control rectangle, (i.e. the diffusing particle contains either more fluorophores or is of higher intensity). Thus, this method will be sensitive to the formation of nascent particles containing higher amounts of eGFP-G3BP1. It is important to note that N&B is not sensitive to immobile aggregates or larger, pre-formed particles.

We monitored N&B of G3BP1 in adult Day 1, 4 and 8 worms with cycles of 30-minute activation of Gαq by carbachol addition and 30-minute recovery by removal of stimulant. Aggregation was assessed by comparing the movement of GFP-G3BP1 expressed in worms to free GFP. Changes in N&B may reflect two major mechanisms; 1 -the formation and dissolution of nascent G3BP1 particles resulting in shifts in points to higher values in N&B plots; 2-G3BP1 and associated proteins absorb onto slow-moving pre-formed particles causing particles to become undetectable by N&B and resulting in little changes in the N&B plots. In **Supplement Fig.1** we show examples and images of Day 1 worms subjected to repeated Gαq stimulation and recovery, with compiled results for single stimulation of Day 1, 4 and 8 in **Table 1** and compiled results for repeated stimulation of Day 1 worms in **Table 2**. The relatively low increase in aggregation indicates a low level of nascent formation of particles during the first stimulation that appears to extend into the recovery phase in Day 1 and 4 worms. Day 8 worms show little response to stimulation or recovery. Repeating the stimulation and recovery cycle in Day 1 worms (**Table 2**) indicate the formation of new particles during the second cycle and this trend reverses in the third suggesting the coalescence of small particles into larger ones that are not detectable by this method.

**Table 1.**
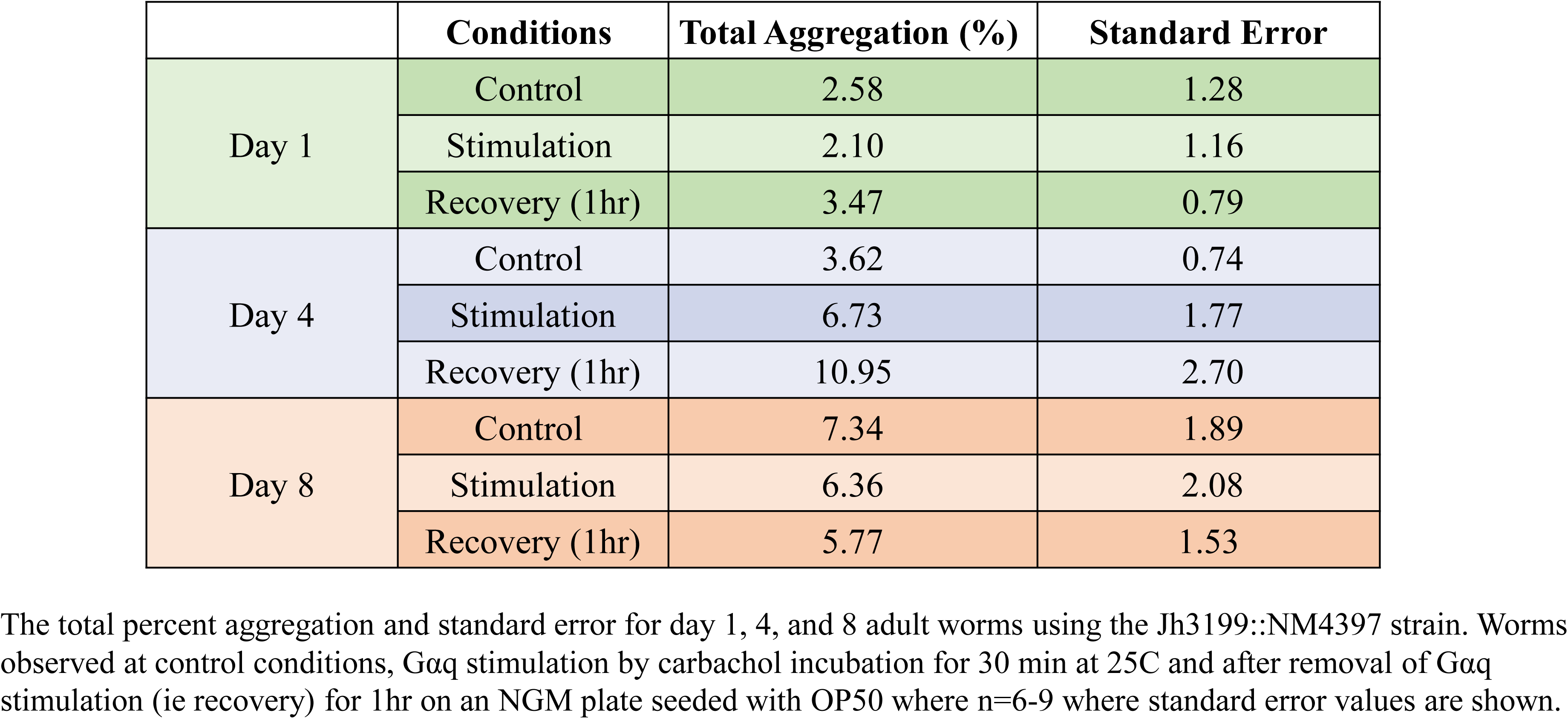
Number and brightness results of G3BP1::gfp in adult C. elegans.

|  | Conditions | Total Aggregation (%) | Standard Error |
| --- | --- | --- | --- |
| Day 1 | Control | 2.58 | 1.28 |
|  | Stimulation | 2.10 | 1.16 |
|  | Recovery (1hr) | 3.47 | 0.79 |
| Day 4 | Control | 3.62 | 0.74 |
|  | Stimulation | 6.73 | 1.77 |
|  | Recovery (1hr) | 10.95 | 2.70 |
| Day 8 | Control | 7.34 | 1.89 |
|  | Stimulation | 6.36 | 2.08 |
|  | Recovery (1hr) | 5.77 | 1.53 |

The total percent aggregation and standard error for day 1, 4, and 8 adult worms using the Jh3199::NM4397 strain. Worms observed at control conditions, Gαq stimulation by carbachol incubation for 30 min at 25C and after removal of Gαq stimulation (ie recovery) for 1hr on an NGM plate seeded with OP50 where n=6-9 where standard error values are shown.

**Table 2.** Number and brightness results for Dai1 G3BP1::gfp C. elegans.

| Conditions | Total Aggregation (%) | Standard Error |
| --- | --- | --- |
| Control | 4.72 | $\pm 3.23$ |
| Stimulation 1 | 5.37 | $\pm 3.09$ |
| Recovery 1 | 4.74 | $\pm 3.22$ |
| Stimulation 2 | 4.85 | $\pm 3.17$ |
| Recovery 2 | 13.10 | $\pm 2.32$ |
| Stimulation 3 | 6.78 | $\pm 4.62$ |
| Recovery 3 | 1.47 | $\pm 0.60$ |

Because N&B can only capture the formation and dissolution of small particles, these studies were followed by observing larger eGFP-G3BP1 particles in mechanosensory neurons by confocal imaging (**Fig. 3**). In Day 1 worms, initial stimulation of Gαq causes little change in the number of stress granules, but a significant increase in their size (**Fig. 3A-B**). After a 30-minute recovery, there is a small increase in the number of particles while the size of the particles remains high, suggesting disassembly of larger aggregates **(Fig 3C-D)**. Repeating Gαq activation further increases the number of particles without significant changes in size, and this number increases after recovery (**Fig. 3A-D**). Stimulating a third time increases the number of particles that then increase in size during recovery. In contrast, Day 4 worms (**Fig. 3E-H**) show a muted behavior where the trends barely reached significance, and Day 8 worms (**Fig. 3 I-L**) show small increases that did not recover. These data show that repeated Gαq activation initiates the formation and growth of stress granules in a robust and reproducible manner in young worms that are reduced in middle-aged worms. Older worms can form stress granules that do not disassemble during the time of the experiment.

**Figure 3.**
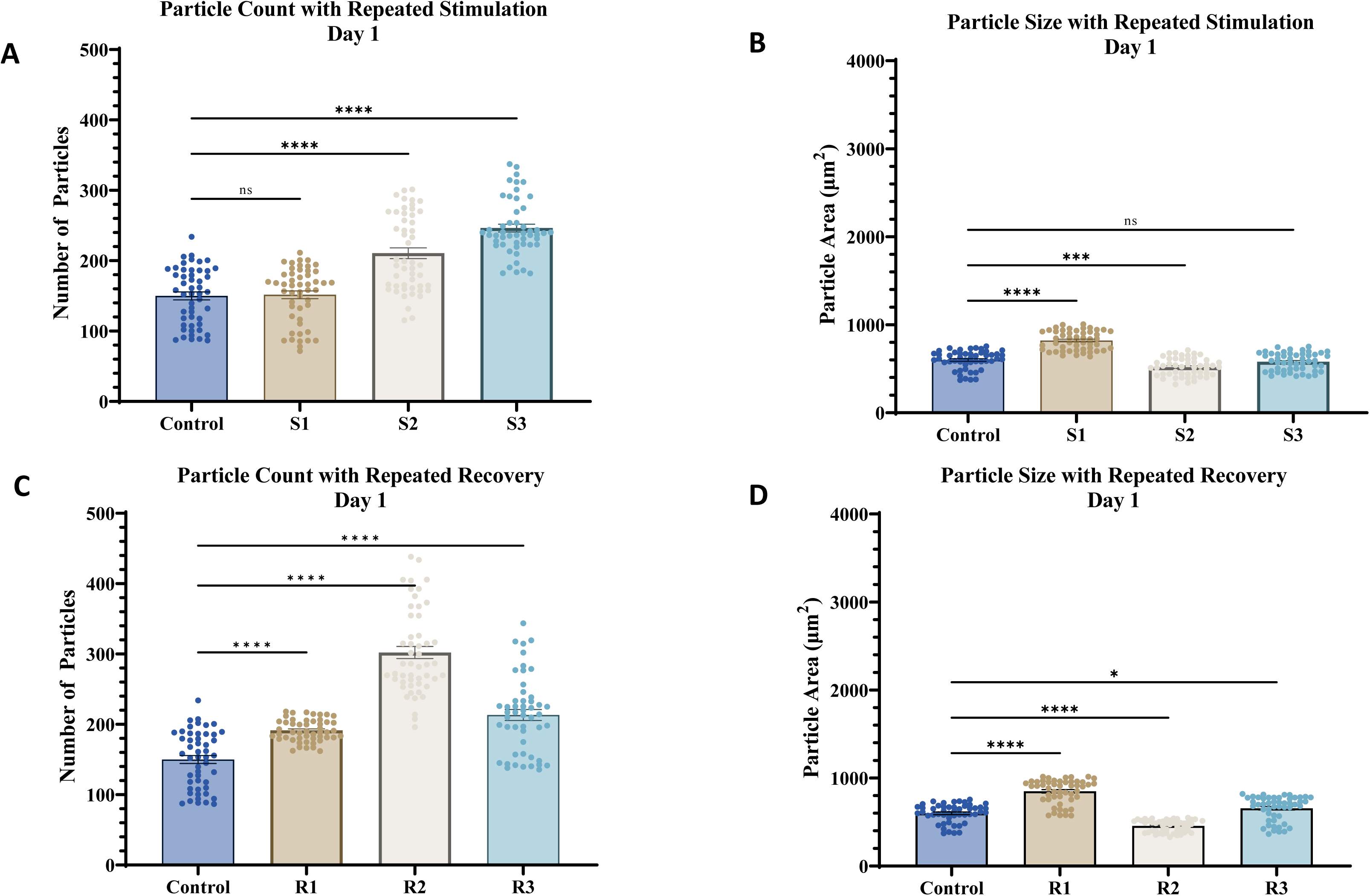
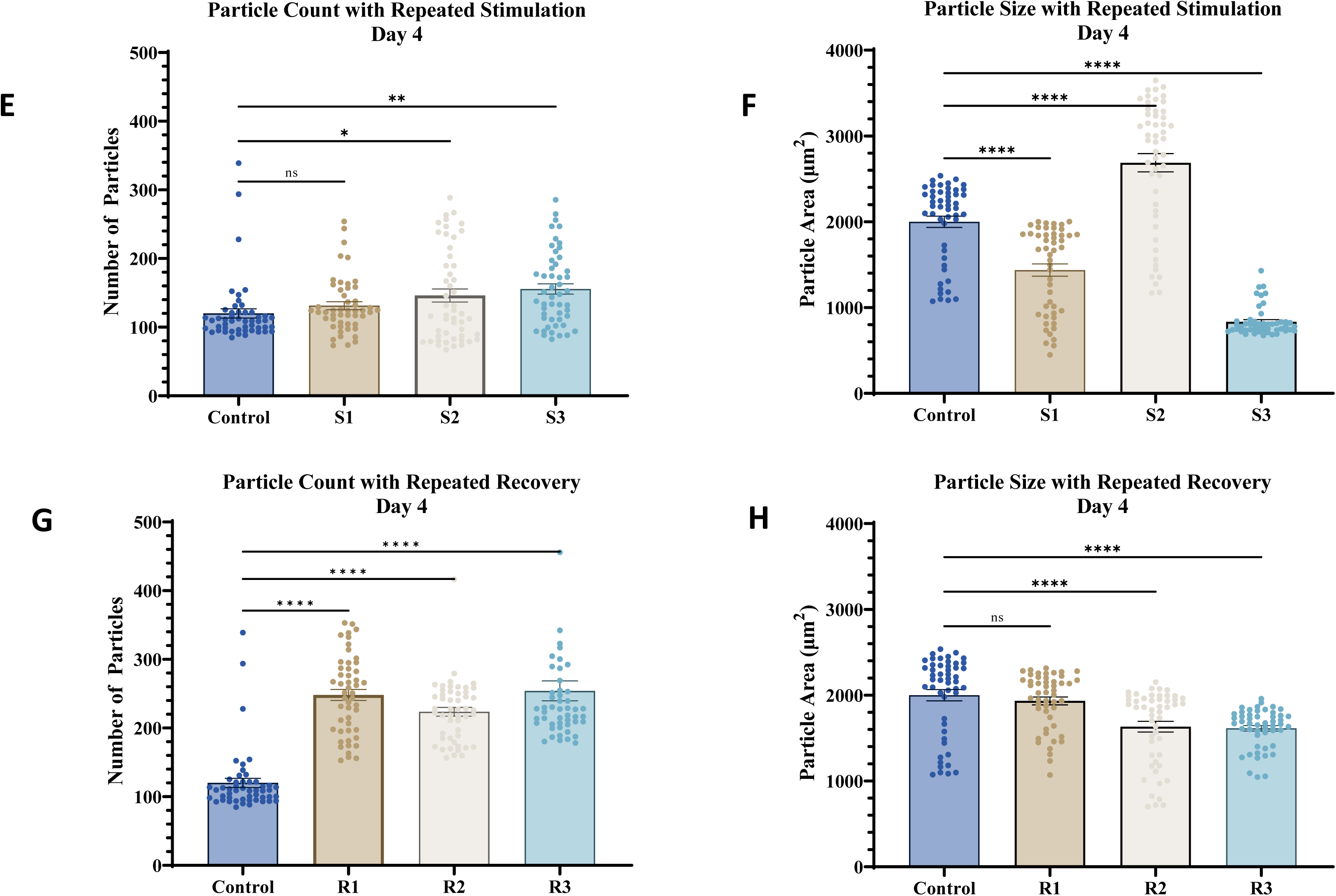
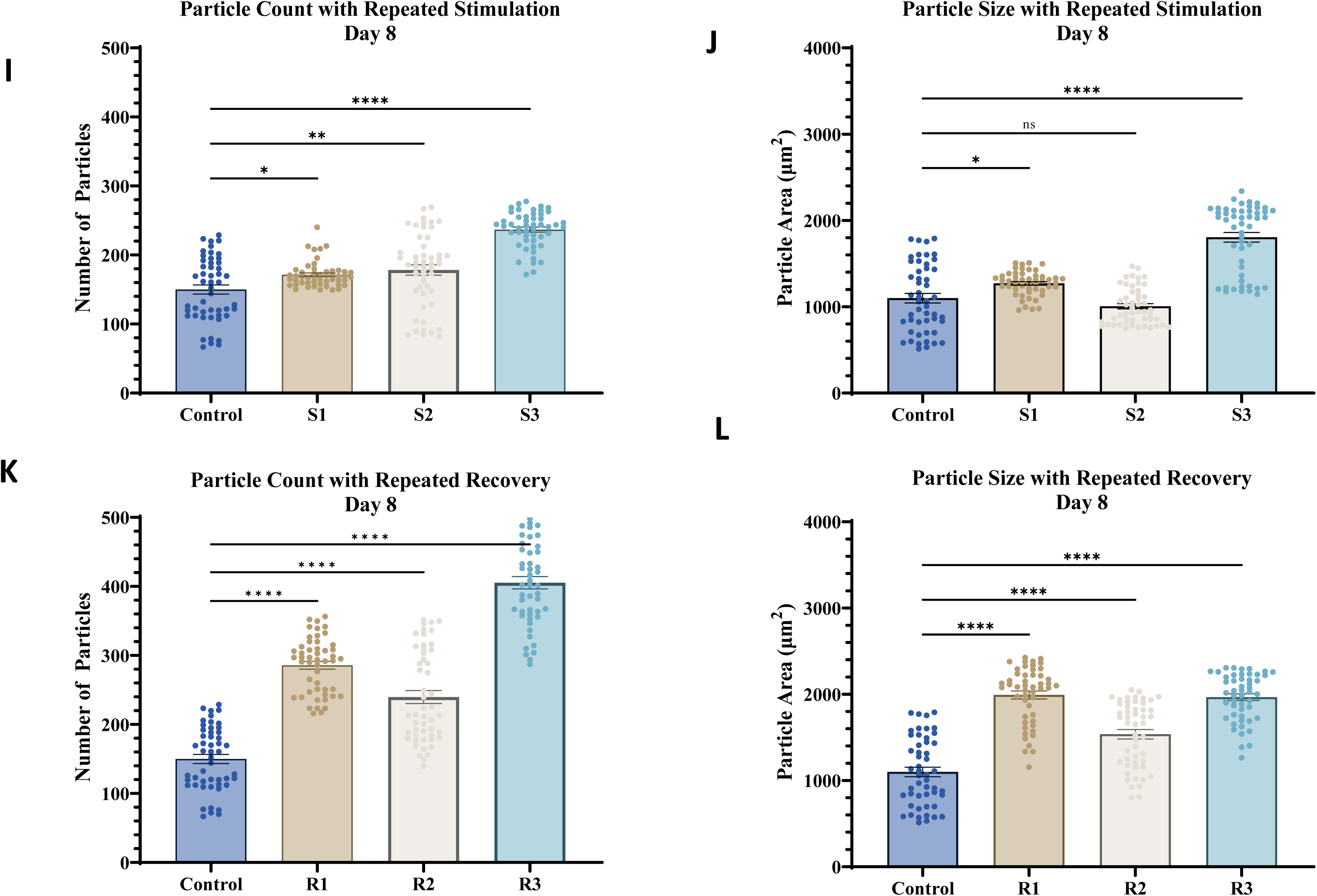
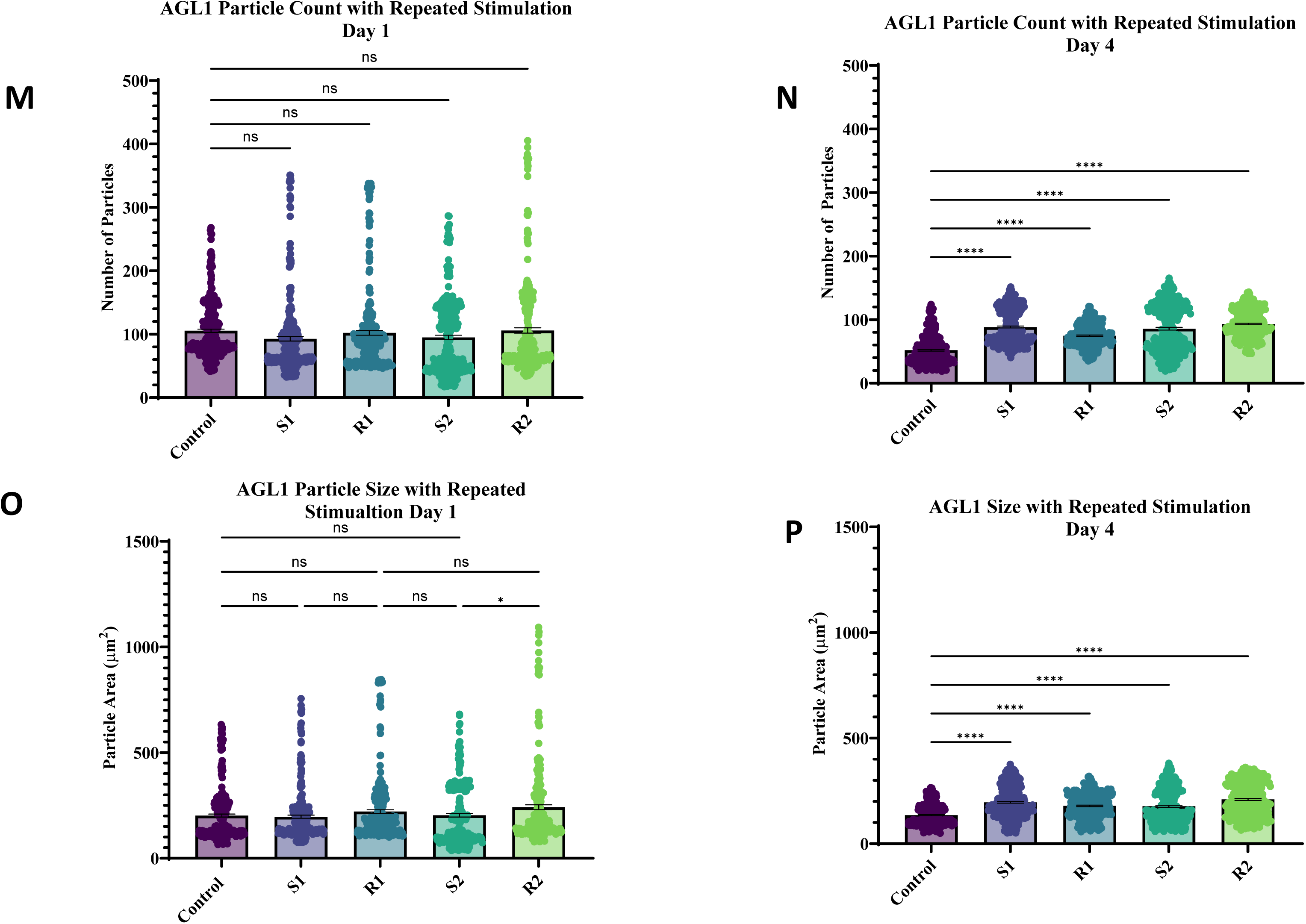
Number and size of stress granules upon repeated Gαq stimulation and recovery. Aggregation of G3BP1::gfp in JH3199::NM4397 *C. elegans* head neurons by confocal imaging using 1 m thick z-stack images were analyzed in adult Day 1 (**A-D**), Day 4 (**E-H**) and Day 8 (**I-L**). G3BP1 particles were observed using confocal imaging under control conditions, immediately following 1 mM carbachol exposure to stimulate Gαq (S1) for 30 minutes at room temperature. After the carbachol exposure was removed and the worms were allowed to recover on a seeded NGM plate (R!). This process was repeated a second (S2 and R2) and third cycle (S3 ad R3). For all ages n= 10-15 and SEM is shown. Data was visualized using GraphPad Prism and analyzed using a One-Way Anova, where p-values are represented on the graph. (“ns” correlates to non-significant data points and **** correlates to a p-value < 0.0001). **(M-O)** Aggregation of ALG-1 in Day 1 and Day 4 adult in transgenic alg-1::gfp PQ530 crossed with transgenic mec-7p::mRFP *C. elegans* where head neurons were imaged confocally with 1m slices to assess analyzed for ALG-1 aggregation on Day 1 (**M-N**) and Day 4 (**O-P**) under control conditions, immediately following stimulation of Gαq at room temperature by carbachol for 30 min (S1), and recovery on a seeded NGM plate for 30 minutes (R1), where n= 10-15 and SEM values are shown. Data were visualized using GraphPad Prism and analyzed using a One-Way Anova, where p-values are represented on the graph. (“ns” correlates to non-significant data points and **** correlates to a p-value < 0.0001).

We compared the formation of G3BP1 stress granules due to Gαq activation in adult Day 4 *C. elegans* with heat stress, whose ability to generate stress granules has been well described (31), **Supplemental Fig 2).** To prevent heat damage (See (32)), we shortened the stress times from 30 to 15 minutes and increased the recovery time to 1 hour. We find that heat shock results in a steady increase in particle number that continues to rise during recovery, and these changes are accompanied by a small increase in particle size. These results show that, in contrast to Gαq activation, subjecting worms to heat shock results in systematic and irreversible increases in the number of stress granules.

### Repeated Gαq stimulation increases the number and size of Ago2 stress granules

Ago2 is the key nuclease component of the RNA-induced silencing complex and will form stalled complexes (i.e. stress granules) when there is imperfect pairing between the Ago2-bound miR and mRNA (33, 34). Our previous studies have shown that under basal conditions, cytosolic PLCβ1 directly binds to Ago2, reducing the ability of Ago2 to form stress granules (18). Here, we followed changes in Ago2 (ALG-1) stress granules in Day 4 adult *C. elegans* tagged with a3xFLAG:GFP for ALG-1 (PQ530) with single and repeated Gαq activation. We find that both the size and number of ALG-1 particles are similar to G3BP1 particles supporting the idea stress granules contain both markers. Repeated stimulation did not significantly change the number of particles in Day 1 worms (**Fig. 3M-N**) but in Day 4 worms, particle number increased slightly and showed a small degree of recovery (**Fig. 3 O-P**). Notably, ALG-1 particle sizes increased with repeated stimulation without completely recovering. However, increases in size were smaller than that seen in G3BP1. Overall, our studies suggest that stress granules associated with ALG-1/mammalian Ago2 accumulate with repeated stress (**Fig. 3M-P**).

### Gαq stimulation protects the level of ATP5f1b

In PC12 cells, activation of Gαq sequesters the *ATP5f1b* gene in stress granules to protect the cell’s energy sources. To determine whether this is also the case for neurosensory neurons in *C.elegans*, we measured changes in the amount of the ATP5f1b protein under control, Gαq activation, and heat stress in Day 1, 4 and 8 worms by comparing changes in immunofluorescence intensity using a monoclonal antibody (**Fig. 4A**). Each age group showed different behavior that reflected the ability of Gαq to assembly and disassemble stress granules (**Fig. 2**). Day 1 worms, which showed larger stress granules that remained large and became more numerous during recovery showed a significant increase with carbachol stimulation and little change with heat. Alternately, Day 4 worms, which showed significant reductions in both size and number with Gαq activation that did not fully recover, showed a decrease in ATP5f1b levels and a slight reduction with heat. Day 8 worms, which showed a decrease in size and number with Gαq stimulation followed by an increase and little change with change in ATP5f1b levels and a decrease with heat. This latter behavior is similar to that seen in PC12 cells that showed protection of ATPsynthase levels with Gαq stimulation in contrast to heat. It is curious that higher levels of ATP5f1b are seen with aging especially in light of the exponential decrease of ATP stores and mitochondrial dysfunction with age (35)(36). It is possible these higher levels are due to compensation for lower ATP levels.

**Figure 4.**
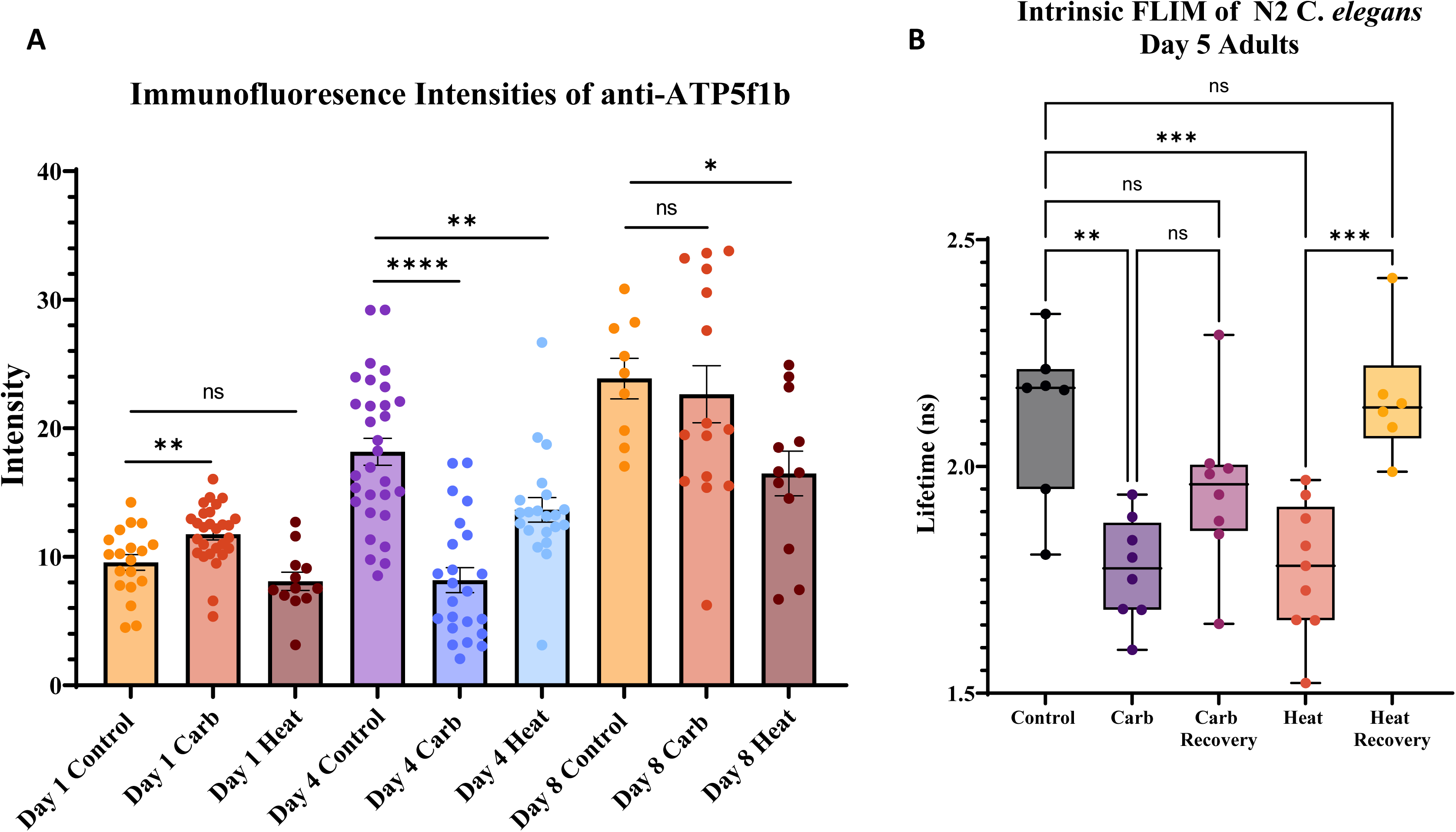
Changes in ATP5f1b levels and redox state with Gaq stimulation and heat shock. **(A)** Wildtype N2 worms were collected at adult Day 1, 4, and 8 stage and analyzed for ATP5f1b levels by immunostaining with a monoclonal antibody under control, Gαq stimulation by treating with 1mM carbachol at room temperature for 30 min, and heat shock at 35°C for 15min. All intensity values were background subtracted and normalized to an unstained control. Student t-tests comparing two groups is given where ** P≤0.001 and ***P≤0.0001. **(B)** Changes in the lifetime of the intrinsic fluorescence of Day 5 worms, which is related to the redox state, with Gαq stimulation by treating with 1mM carbachol at room temperature for 30 min, and heat shock at 35°C for 15min where both were followed by a 30 min recovery.

We assessed changes in the redox state of the whole organism by monitoring changes in intrinsic fluorescence. In these measurements, the lifetimes associated with the intrinsic fluorescence of NAD(P)+/NAD(P)H and, to a lesser extent FADH2, report on cell health (37, 38). We find that Gαq stimulation reduces the overall lifetime, lowering the redox state corresponding to increased oxidation (**Fig. 4B**), and recovers after the stimulus is withdrawn. This same behavior is seen with heat shock. These data show that the redox state of Day 5 worms are responsive to stress and fully recover when the stress is withdrawn.

### Aging impacts the time-dependence of Gαq stimulation changes in locomotion

In previous studies, we found that Gαq stimulation causes contraction of *C.elegans* neurons that disrupts neural connections and locomotion (14). In cultured cells, retraction can only be reversed by the addition of nerve growth factor, but in Day 5 *C*.*elegans*, reversal occurs naturally after the stimulus is removed. Because Gαq stimulation disrupts neuronal connections, we studied the effect of repeated Gαq stimulation on movement by assessing *C.elegans* locomotion.

Changes in *C.elegans* peristaltic speed (39) after carbachol addition at various time points were measured in Day 1, 4 and 8 worms using the crossed strain (JH3199::NM4397) (**Fig. 5A-C**). We find that Day 1 worms show an initial increase from 180-200 μm/s after the addition of carbachol that reaches a maximum average of 600 μm/s after 15-minute post-Gαq stimulation. In contrast, Day 4 worms showed an initial drop in speed in the first 30-minute post stimulation and an average maximum of ∼500 μm/s, that was not reached until 60-minute post stimulation. Like Day 1 worms, the speed of Day 4 worms recovered after their maximum speed was obtained. We note that while the maximum peristaltic speeds of Day 1 and 4 were similar, their onset times were very different. Even more different were Day 8 worms, which showed a small response to stimulation, but their speeds were much slower (200-360 μm/s) in accord with previous studies showing functional decline in motor activity (40).

**Figure 5.**
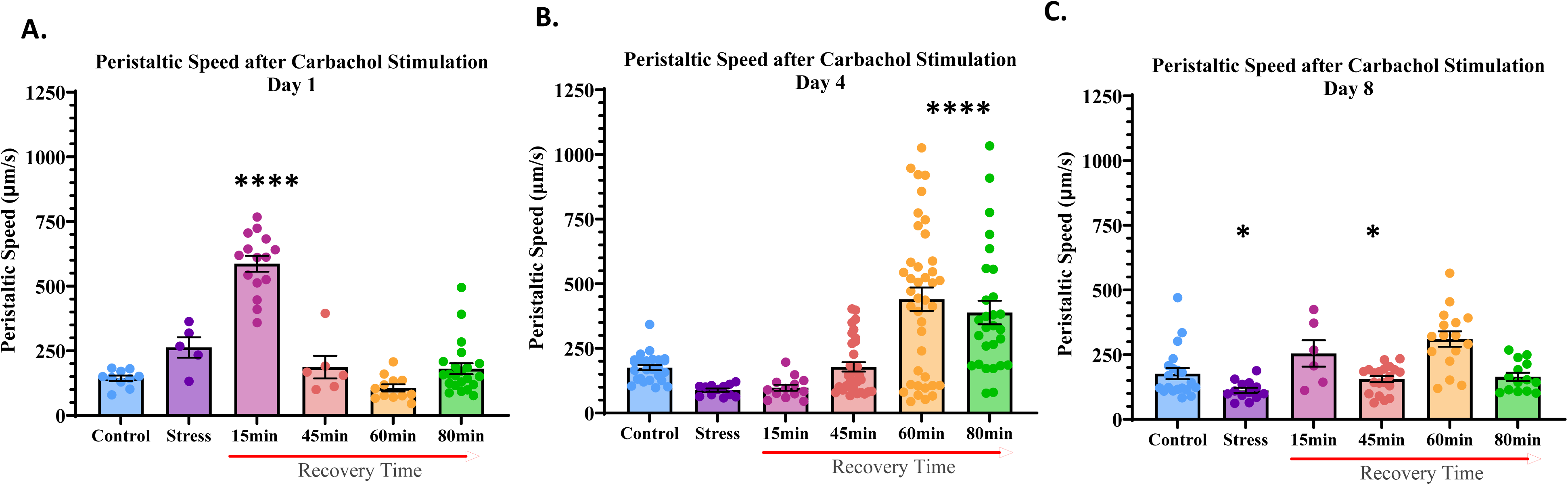
Recovery of peristaltic speed of JH3199 *C. elegans* after Gαq stimulation. Worms were stimulated with 1mM Gαq at room temperature and seeded on a NGM seeded plate at room temperature followed for 15-80min in Day 1 **(A)** n= 6-22, Day 4 **(B)** n=6-20, and Day 8 **(C)** n=12-49 adults. Worms were stimulated using 1mM carbachol for 30min at room temperature. Peristaltic speed (Peristaltic Speed (µm/s)= [forward track length + reverse track length](24) / time (24)) was recorded using an inverted camera and software from MBFBioscience. Videos were recorded for 1min and analyzed using WormLab camera software. Data was visualized using GraphPad Prism and analyzed.

### Repeated Gαq stimulation alters neuron morphology

The onset of increased peristaltic speed may be due to the ability of the synapses to recover after Gαq stimulation (14). To support this idea, we imaged changes in the morphology of *C.elegans* mechanosensory neurons in the head, focusing on the nerve ring and dendritic spine structures with single and repeated Gαq activation (**Supplemental Fig. 3**). In (**Fig. 6A),** we plot the total number of morphological changes (i.e. beading and synapse disruption), as seen using a 63x objective (**Fig. 6C**). Under control conditions, neurons in Day 4 worms do not show abnormal morphology but show extensive changes with stimulation that reverse with recovery. This same behavior is seen during the second cycle of stimulation and recovery and is consistent with the initial reduction in locomotion and followed increased locomotion. Alternately, Day 8 worms show a significant number of morphological abnormalities under control conditions that decrease with Gαq stimulation, and this trend reverses during recovery, with similar trends in the second application of stimulation and recovery. When beads and nerve ring disruptions are viewed separately (**Fig. 6B**), we find that responses in nerve ring morphology are minor compared to changes in beading, which dominate the responses. These changes in morphology are interpreted to reflect contraction of the underlying muscle with Gαq stimulation.

**Figure 6.**
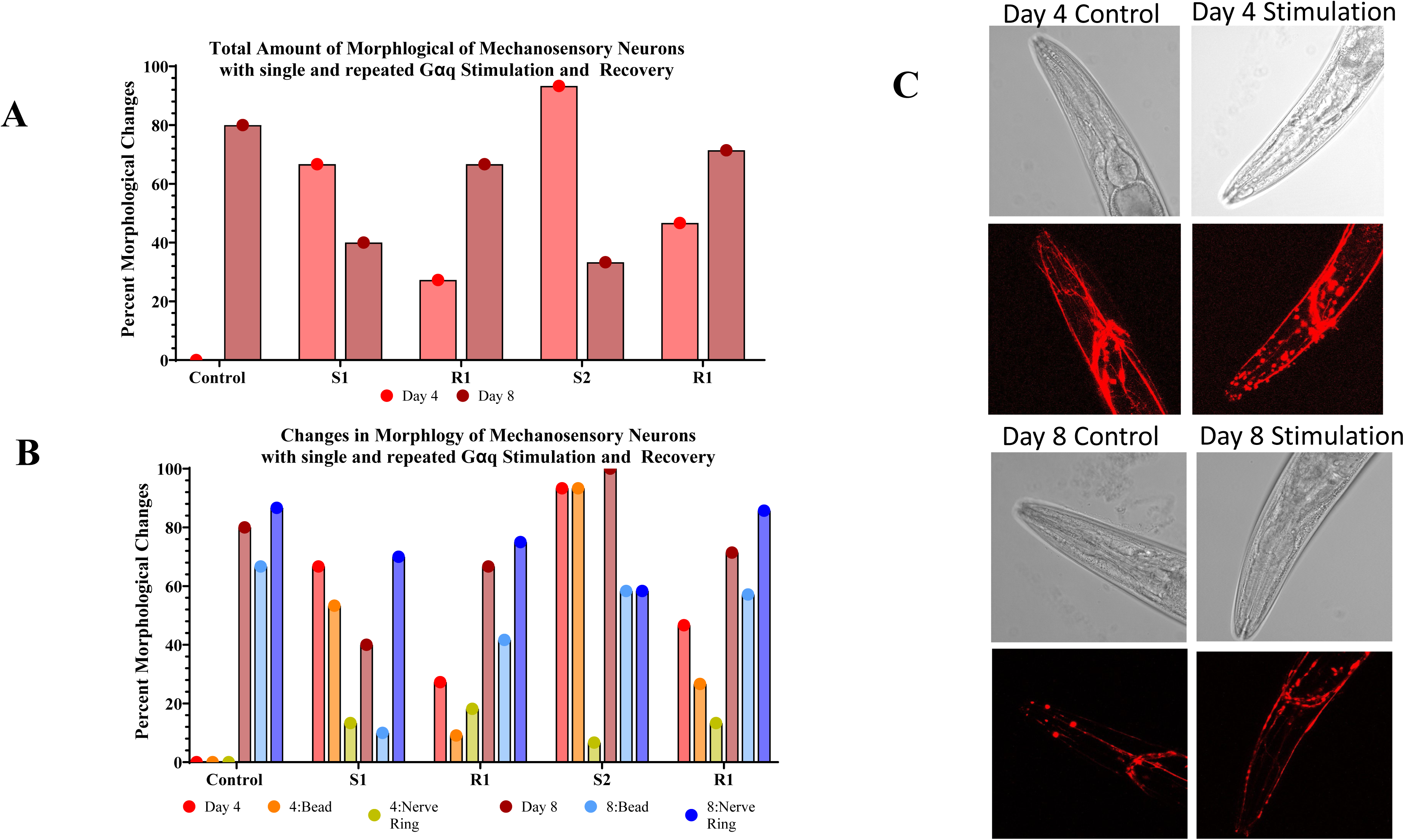
Percent of normal morphology in mechanosensory nerves in NM4397 upon Gαq stimulation and recovery. Total changes (**A**) and nerve ring rupture and dendritic spine beading (B) were compiled by confocal images (**C-E**) under control conditions, one round of Gαq stimulation by 1mM carbachol at room temperature (S1), and recovery at room temperature on an OP50 seeded NGM plate (R1), followed by a second another cycle (S2 and R2). To visualize morphology throughout the worms head region, z-stack imaging was utilized which captured 1um thick slices throughout the worms head. For all conditions and ages n=11-15. All images were processed using Fiji software and data was processed using GraphPad Prism.

## DISCUSSION

This study focuses on the role of repeated Gαq stimulation on the health of *C.elegans*. Stimulation of Gαq (EGL-30), whose gene has ∼80% homology to mammalian Gαq (41), is a normal physiological event that is positively associated memory through calcium-mediated activation of CREB (42), and is associated with movement and *C. elegans* locomotion (see (43)). Recent studies (44) found that expressing constitutively active Gαq (EGL-30) preserves long term associative memory in older worms. Interestingly, this mutation does not impact locomotion, suggesting that its impact on worm health is not simply due to elevated calcium levels, and that other mechanisms may be involved. In contrast to constitutively active EGL-30, *C.elegans* with other mutations in EGL-30 have egg-laying defects, reduced viability, pharyngeal pumping, and movement (see (45)). Together, these studies suggest that continuous Gαq activity has positive effects, presumably due to adaptive mechanisms, whereas defects in this pathway are detrimental. However, in these studies, we find a decrease in worm lifespan when Gαq is stimulated once a day, noting that recovery of locomotion from calcium-induced Gαq signals and the metabolic state recover quickly (**Figs. 5-6**).

To understand the effects of single and repeated Gαq stimulation on the neuronal health of *C.elegans,* we first examined the impact of Gαq-mediated stress granule formation (18). Stress granules can be formed by different environmental perturbations and those assembled by Gαq activation are expected to have different compositions and properties than those formed under other treatments. For example, Ching and coworkers found in *C. elegans* that dietary restriction mediates stress granule formation through a pathway distinct from heat shock and starvation (46). We reasoned that repeated Gαq may impact worm lifespan, in part, through the formation of stress granules. This idea is based in cell culture studies (18) and *C.elegans* studies showing that intracellular aggregates, which may include stably assembled stress granules, are found in many neurodegenerative disease (47).

Previously, David and coworkers followed protein aggregates in *C. elegans* as a function of age by characterizing the amount and content of insoluble protein aggregates (8). Here, we followed the number and size of G3BP1 stress granules using the canonical stress granule marker G3BP1 in *C. elegans* from Day 1 to 15. We delineated changes in stress granules in terms of number which corresponds to assembly and disassembly of particles, and size which corresponds to the association or dissociation of small G3BP1 to larger complexes, such as stalled ribosomes. In contrast to observing an increase in protein aggregation with age, we find that the number of G3BP1 stress granules remains constant throughout the worm’s lifetime and decreases somewhat in size. These results suggest that the aggregates seen in previous studies (8) were not true stress granules but other complexes formed from dysfunction in protein degradation, or irreversible entanglement in disordered regions of proteins that result from reduced proteosome activity (48).

Compared to undifferentiated PC12 cells, *C.elegans* mechanoneurons contain ∼10 fold less particles that are ∼10 fold larger in size under basal conditions. One possible reason for this difference is that stress granules in undifferentiated cells contain a higher percentage of miRs, while stress granules in differentiated cells contain higher levels of mRNA. *C.elegans* stress granules change in size with Gαq activation rather than both size and number as is seen in cultured cells as the neurons respond to activation by sequestering different transcripts. Additionally, the behaviors depend on the age of the worm with little or no changes in younger and middle-aged worms but reductions in particle size in older worms that do not recover from Gαq activation. These changes in particle size may reflect absorption / desorption of small protein oligomers as the cells respond to stimulation. Support for this idea comes from the lack of response in G3BP1 oligomerization in Day 4 and Day 8 worms as seen by N&B (18, 30, 49) (**Table 1**).

Although Gαq stimulation does not greatly impact stress granule size and number, we find that repeated stimulation results in a steady increase in particle number and size suggesting the formation of new stress granules as well as the growth of pre-formed stress granules. However, recovery to basal levels was incomplete, suggesting that stress granules may accumulate with repeated stimulation, and the propensity for accumulation appears to increase in older worms (**Fig. 3**). We note that selected specific conditions of stimulation and recovery that were experimentally straightforward and most reproducible experimentally, but these conditions were sufficient to show the possibility of stress granule accumulation. However, it is unclear that these accumulated complexes underlie neurodegenerative disorders (see (24, 50)). Comparing stress granule behavior in response to repeated Gαq stimulation with heat shock shows that both conditions result in stress granule accumulation. With heat shock, the composition of these accumulated complexes is probably less specific compared to the more regulated Gαq-induced stress granules.

While G3BP1 is the *de facto* stress granule marker, Ago2 is a binding partner of PLCβ and is released as PLCβ binds to Gαq upon stimulation (18, 26). We followed the size and number of ALG-1, the worm equivalent of Ago2, with single and repeated Gαq stimulation whose trends were similar, but smaller than those seen with G3BP1 consistent the idea that ALG-1 stress granules are a subset of G3BP1 stress granules (18). In cultured cells, Gαq activation promotes the formation of Ago2 stress granules that protect and sequester two RNAs, *Chgb* and *ATP5f1b* (see (26)). We tested whether this is the case in *C.elegans* neurons. Monitoring ATP5f1b levels by immunofluorescence in Day 1, 4, and 8 worms show age-dependent changes that mirror their stress granules assembly. It is unclear why younger worms protect and enhance levels while middle-aged worms do not, and our data suggest it may be related to the size and number of Gaq-induced stress granules. Nevertheless, these results support the idea that Gαq stimulation induces stress granules that sequester specific RNAs that include *ATP5f1b.* We note that immunostaining of ATP5f1b in Day 8 worms was more intense than Day 1 and 4. We believe these higher intensities are due to better accessibility of the antibody to the epitope due to loss of mitochondrial integrity rather than higher expression levels in Day 8 worms (28, 36). Efforts to determine levels by western blotting were unsuccessful.

Besides impacting the assembly and disassembly of stress granules, Gαq stimulation results in morphological changes in *C.elegans* neurons that may be directly traced to neurite retraction caused by calcium-dependent cytoskeletal changes, PIP_2_ hydrolysis, and endocytosis of ligand-bound Gαq coupled receptors (14). Retraction of neurites will disrupt weaker synapses causing cessation of locomotion. Wolkow and coworkers have found that locomotion of *C.elegans* declines with age, and that this decline can be rescued by muscarinic agonists (9). Here we find that the onset and amount of increased speed is age dependent, with Day 1 worms displaying a short delay and a large increase in speed, and Day 8 worms show little change (**Fig. 5**). Because these age-dependent behaviors may reflect disruption of synapses upon Gαq stimulation and subsequent recovery, we characterized changes in morphology of mechanosensory neurons focusing on nerve ring disruptions and neural beading. Gαq stimulation of Day 4 worms causes a small amount of nerve ring disruptions that remain fairly constant with repeated stimulation and recovery. Gαq stimulation causes a large amount of beading that is reduced with recovery and increases and recovers again with a second stimulation / recovery cycle. Day 8 worms display a large amount of nerve ring disruption even before stimulation which remains high while the large amount of beading is reduced during the first stimulation and remains high after. These results suggest that beading is the main morphological change associated with Gαq stimulation and recovery. Differences between nerve ring disruption and beading may reflect contraction of neural extensions due to contraction of the surrounding muscle (51). This contraction is expected to help the recovery of retracted synapses. While middle-aged worms respond and recover quickly from these physical changes, older worms do not. These worms have impaired nerve morphology and show diverse and broad responses most likely due to their lack of Gαq response and muscle contraction. These studies highlight the robust stimulation of middle-aged worms as compared to older ones.

In this study, we show that repetitive Gαq stimulation reduces *C.elegans* lifespan though mechanisms that may include stress granule accumulation and generation of abnormal neuron morphology in an age-dependent manner. While it is doubtful that constitutively active Gαq could rescue these effects, it is possible that increasing Gαq and/or PLCβ levels would help disassembly stress granules and promote recovery.

## MATERIAL AND METHODS

### *C. elegans* strains and maintenance

All strains were obtained from *Caenorhabditis* Genetics Center. Strains, JH3199 (gtbp-1, (as2055[gtbp-1::GFP])); NM4397 (jsls973[mec-7p::mRFP + unc-119(+)]); and PQ530 (alg-1, (ap423[3xflag::gfp::alg-1])). Standard culture methods were used to maintain C.elegans (52). All strains were grown on nematode growth media (NGM) agar plates seeded with OP50 *Escherichia coli* (*E. coli*). The strains were maintained at 20°C.

### *C. eleg*ans Crossing Procedure

To take advantage of multiple fluorescent markers at a time, *C.elegans* strains were crossed using a mild heat-shock procedure. Strains of interest were placed on NGM OP50 plates as L1 larvae. Plates were the mildly heat shocked in either an incubator (30°C) or placed inside down in water bath (30°C) for 4-6 hours. Heat-shocked L1 larvae plates were then stored at 20°C until L4 progression. Once L4 stage was reached, all plates were observed for the presence of male *C.elegans*. Once males were identified for one strain, one male was placed with 5 hermaphrodites from another strain of interest. Once progeny formed, crossed *C. elegans* were confirmed using confocal microscopy taking advantage of each strain’s unique fluorescent marker. This procedure was followed to cross JH3199 with strain NM4397 (JH3199::NM4397) and strain PQ530 with strain NM4397 (PQ530::NM43967).

### Synchronization assay

Worms were synchronized to observe age-related functions using a typical bleaching protocol (53).Gravid worms were collected off NGM OP50 plates, washed with M9 Buffer (3g KH_2_PO_4_, 6g Na_2_HPO_4_, 5g NaCl, 1mL 1M MgSO_4_ to 1L H20, sterilized by autoclave), and bleached to release eggs. Bleaching solution was made fresh (diH_2_0, 5% NaOCl (35% final volume), and 5N NaOH (20% final volume)). Once released, eggs were spun down and washed with M9 buffer. Eggs and bleached gravid adults were plated on fresh NGM OP50 plates and allowed to progress to L4 larvae. Once at an appropriate age, synchronized worms were moved to fresh OP50 plates and analyzed for various assays.

### Application of stress conditions

Carbachol was used to stimulate the Gαq subunit to exhibit stress. Carbachol in powdered form was obtained from Sigma Aldrich (carbamoylcholine chloride, Catalog #C4382) and dissolved in water to a final concentration of l mM at a volume of 25mL. 1mM carbachol was aliquoted into 1mL portions to be used for each experiment. The solutions were kept at -20°C for storage. For single carbachol exposure, worms were washed with M9 buffer and placed into a 15mL centrifuge tube. Worms were allowed to sink to the bottom and M9 buffer was removed. 1mL of 1mM was added to the centrifuge tube for 30min at room temperature. Once stress was complete, carbachol was removed and worms were washed in M9 buffer and moved to a fresh OP50 plate for recovery periods at room temperature. For repeated carbachol stimulation, this process was repeated with the same *C. elegans* for two or three times with recovery periods in between. For heat shock, worms on an NGM OP50 plate were placed upside down in an oven at 35°C for 15 minutes.

### Locomotion assay

To access locomotion patterns, worms were collected and analyzed using WormLab® and analyzed using WormLab® Software (MBF Bioscience, Williston, VT). Worms were placed on NGM dishes with or without OP50 depending on the condition. After 30sec of adaptation, worms locomotion was recorded using a Basler acA2500-14um USB 3.0 camera with the ON Semiconductor MT9P031 CMOS sensor and a resolution of 5MP resolution at 14 frames/s. Camera settings were adjusted to capture all worms on the plate. Wormlab® is a software that enables tracking and image analysis to collect animal data from videos to enable data automation. Each plate was recorded for 1-5min at room temperature and exported as an image sequence. To analyze, each sequence was converted to a project by setting the plate width, thresholding, and manually selecting worms for the analysis to obtain locomotion data. Each plate was analyzed for peristaltic speed in um/s and each worms speed was recorded. Data was exported at a .csv files and further manipulated in Microsoft Excel and GraphPad Prism software.

### Lifespan assay

Worms were assessed for lifespan measurements using a manual counting assay. Strains of interest were synchronized (See Synchronization Assay) and all released eggs were allowed to grow to the L4 stage. Once at L4, a set number of worms (25-50) were moved to a fresh NGM OP50 plate. After 2 hours on that plate, manual counting began at Day 1 adult stage. Worms were moved over to fresh OP50 plates every day for the first week until adults reached Day 6 and no longer laid eggs to ensure the correct generation was analyzed. From Day 6, worms were then moved every other day and manually counted until lifespan was complete. Dead worms were identified with the lack of response to gentle prodding of a platinum wire near their head region. Dead worms were removed from the plate to ensure accurate count.

### Preparation for fluorescent microscopy

To image, worms were placed onto an agar pad using 2% noble agar on a cover slip with a paralytic agent for viewing. To paralyze, worms were placed in 0.5uL of 1mM tetramisole hydrochloride (Sigma Aldrich, Catalog #L9756) prepared in diH_2_0. For each 0.5uL drop, 2-8 worms were placed before the drop was dry. Once worms were placed on the agar pad, another cover slip was placed on top to ensure worms stayed hydrated during imaging.

### Particle analysis measurements

Worms and cells were imaged using a 2-photon MaiTai laser (Spectra-Physics) (excitation 800 nm at 80 MHz) and a Nikon inverted confocal microscope in an ISS Alba System. Worms were imaged using a 60x water objective to microscopically count the number of particles per μm^2^ formed under different conditions. For each condition, 10-15 worms were prepared on agar slides (See Preparation for Fluorescent Microscopy) and z-stack measurements were taken at 1.0 μm per frame of the worm’s head region and analyzed for particles. All z-stack measurements were combined to generate a three-dimensional image for each sample before analyzing the number of particles per sample and averaging the results. Results were portrayed using GraphPad Prism software and statistical significance was calculated using GraphPad Prism with a one-way Anova.

### Number and brightness measurements

Number and brightness (N&B) measurements and theory are fully described previously. Images were collected using a 2-photon MaiTai laser (Spectra-Physics) (excitation 800 nm at 80 MHz) and a Nikon inverted confocal microscope in an ISS Alba System with a 60x water objective. All images were collected in the RICS and N&B element of VistaVision software. Experimentally, we collected 200 worm images in which we viewed strain JH3199 (gtbp-1::GFP) at a rate of 4 μs per pixel. The images obtained for each worm were 256×256 pixels. N&B data were analyzed with SimFCS software (www.lfd.uci.edu) and regions of interest (256 × 256 box) were analyzed from a 320 × 320-pixel image where the size of the SimFCS4 boxes used was determined by the amount of monomeric protein seen in unstressed control worms. Any pixels on the B vs I plot that were outside of this region were determined to be protein aggregation due to the stress applied. Percent aggregation was calculated based off of total number of pixels outside of the control region, divided by the total number of pixels in the cell. The percent aggregation corresponding to either oligomerization or aggregated monomeric protein are also reported using text in the respective color.

### N&B analysis

N&B defines the number of photons associated with a diffusing species by analyzing the variation of the fluorescence intensity in each pixel in the cell image. In this analysis, the apparent brightness [B], in each pixel is defined as the ratio of the variance, σ, over the average fluorescence intensity <K>.

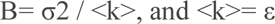

where n is the number of fluorophores. The determination of the variance in each pixel is obtained by rescanning the cell image ∼100 times as described earlier. The average fluorescence intensity, <K>, is directly related to the molecular brightness, ε, in units of photons per second per molecule, and n. B can also be expressed as

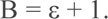

The apparent number of molecules, N, is defined as N = ε / (ε + 1). Statistical Analysis. Data were analyzed with the Sigma Plot 14.5 and GraphPad Prism 9 statistical packages, which included the student’s t-test and one-way analysis of variance (ANOVA).

### Confocal imaging

Confocal images of worms mounted on #1.5 coverslips were obtained using a Zeiss LSM510 Meta inverted confocal microscope. Worms head regions were captured using a x40 (water objective) and acquired using z-stacks with 1micron thick slices to capture the whole head region. Images were analyzed using ImageJ software as a z-project which stacks all slices at the average fluorescence intensity and images were saved as a singular frame.

### Fluorescence lifetime imaging measurements (FLIM)

Phase modulation FLIM measurements were performed on the dual-channel confocal fast FLIM (Alba version 5, ISS Inc.) using a two-photon titanium-sapphire laser and a Nikon Eclipse Ti-U inverted microscope as described (15). The lifetime of the laser was calibrated each time before experiments by measuring the lifetime of Atto 425 in water with a lifetime of 3.61 ns at 80 MHz. Worms were excited at 740 nm for intrinsic measurements, and emission spectra were collected through a 525/50 bandpass filter. For each measurement, the data were acquired using FastFLIM mode on VistaVision software.

### Immunohistochemistry

For immunohistochemistry, a procedure was adopted from previously established method (See 54, 55). *C.elegans* were first synchronized using the synchronization protocol (see above) and collected at Day 1, Day 4, and Day 8. When at the appropriate age, 70-100 transgenic worms were collected with 1mL of M9 buffer, spun down, and M9 was removed. Worm pellets were washed 3 times to remove any bacteria. 1mL of cold boiling extraction buffer was added to the worm pellet (20mM potassium phosphate, pH 7.4, 2mM EDTA, 1% Triton-x-100, and protease inhibitors (ThermoFisher Scientific cat#87786). Worms were boiled for 5min at 95°C and vortexed for 5 seconds every minute. Extraction buffer was removed, and worms were blocked in 10% fetal bovine serum in prepared antibody buffer (700mL distilled water, 100mL 10X PBS, 5mL Triton X-100, 2mL 0.5M EDTA (pH 8.0), 1g BSA, 5mL 10% sodium azide solution, pH 7.2 and brought to final volume of 1L with distilled water) for 1 hour. Worms were probed with primary ATP5f1b (Abcam ab#177991) prepared in antibody buffer at a 1:500 dilution overnight at 4°C. Worms were washed x3 with DPBS and spun down between washes. Worms were then probed with Alexa Fluor™ 488nm goat anti-mouse (ThermoFisher Scientific cat#A11001, Lot#1572559) prepared in antibody buffer at a 1:2000 dilution for 2 hours. Worms were washed 3 times with DPBS and mounted on untreated microscope slides with 10ul of DPBS. Once on slide, worms were allowed to settle for 1-2min and then covered with a 1.5mm coverslip and frozen in dry ice. Slides were kept on dry ice for 5min then sealed for confocal microscopy.

## Acknowledgements

The authors would like to thank Dr. Lela Jackson (Agilent) for her initial help, and Caroline Muirhead and Elizabeth DiLoreto and Dr. Jagan Srinivasan lab (Dept of Biology & Biotechnology WPI). This work was supported by funding from the Richard Whitcomb foundation.

## Author Contributions

MR carried out all of the experimental studies, analyzed results and helped write the manuscript. SS designed the studies, helped analyze the results and wrote the manuscript.

## Data Availability

All data are available upon request.

## SUPPLEMENTAL DATA

**Supplemental.**
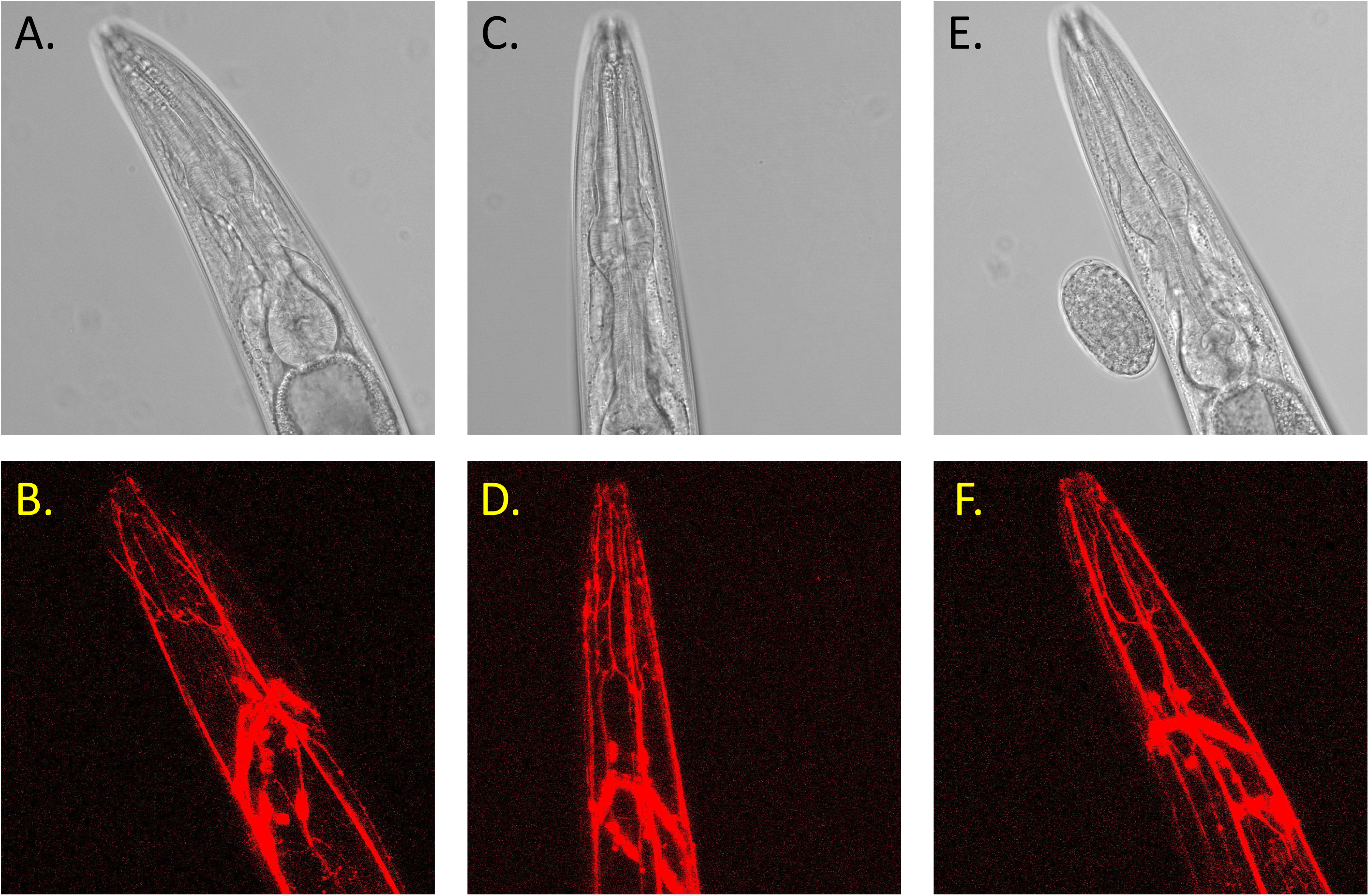
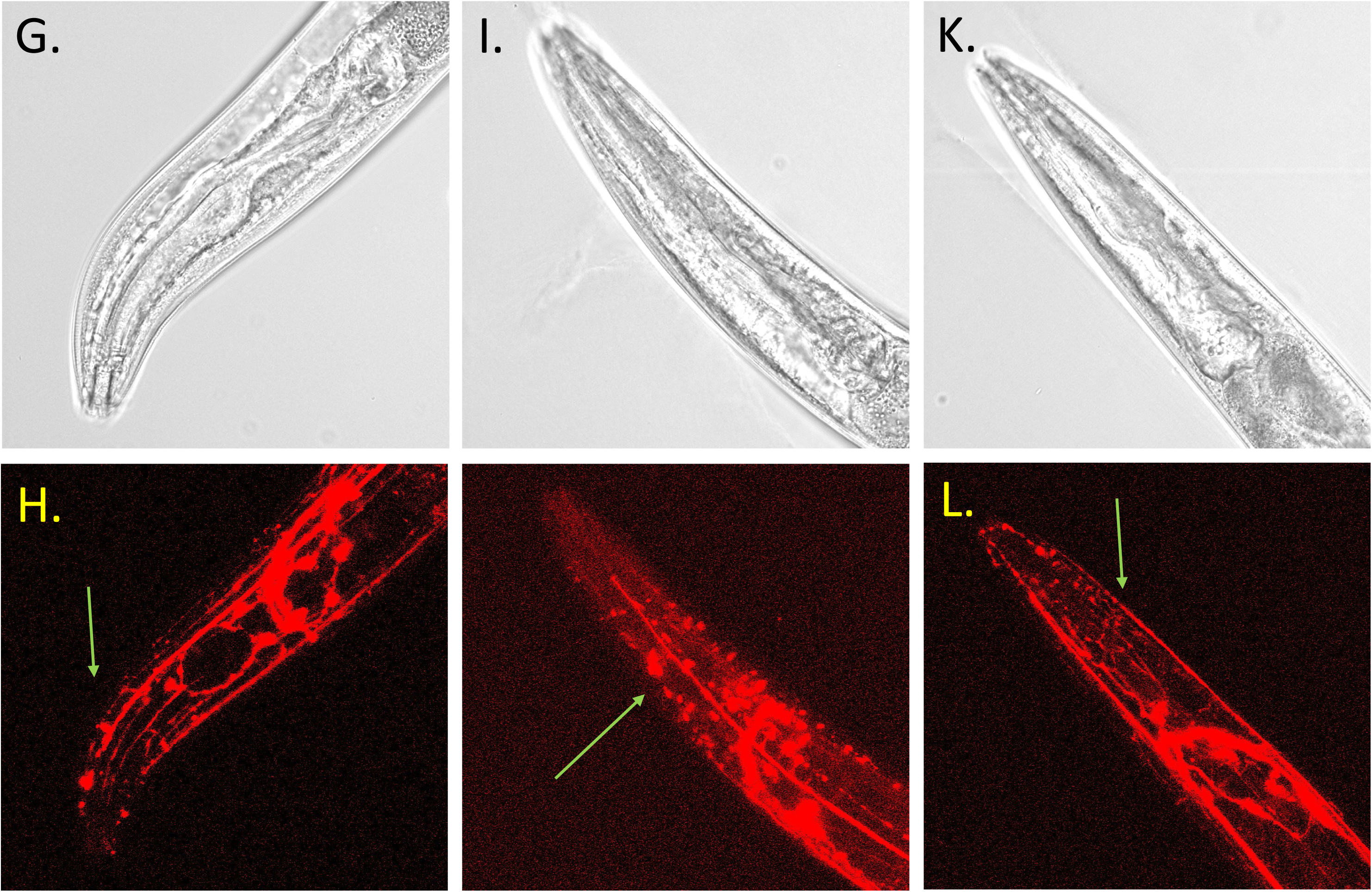
Morphological changes of *C. elegan* head neurons upon repeated Gαq stimulation. Using the NM4397 strain the mechanosensory head neurons were labeled with RFP (mec-7p::mRFP). *C. elegans* were grown on NGM OP50 plates until day 4 of adult hood. Day 4 adults were stimulated with 1mM carbachol for 30min at room temperature to stimulate Gαq. Following carbachol exposure, worms were immediately paralyzed and prepared for confocal imaging (See Methods). Worms were imaged using a 60x water objective at control **(A-F)** conditions and upon Gαq stimulation **(G-L)**. Worms were excited at 543nm and bright field and fluorescent images were collected. The green arrows indicate the beading observed in the head region. To visualize morphology throughout the worms head region, z-stack imaging was utilized which captured 1um thick slices throughout the worms head. For both conditions n=11-15). All images were processed using FiJi software.

## REFERENCES

1. Tissenbaum HA. Using C. elegans for aging research. Invertebrate Reproduction & Development. 2015;59(sup1):59–63.

2. Uno M, Nishida E. Lifespan-regulating genes in C. elegans. NPJ Aging Mech Dis. 2016;2:16010.

3. Antebi A. Genetics of Aging in Caenorhabditis elegans. PLOS Genetics. 2007;3(9):e129.

4. Suriyalaksh M, Raimondi C, Mains A, Segonds-Pichon A, Mukhtar S, Murdoch S, et al. Gene regulatory network inference in long-lived C. elegans reveals modular properties that are predictive of novel aging genes. (2589-0042 (Electronic)).

5. Ross CA, Poirier MA. Protein aggregation and neurodegenerative disease. Nat Med. 2004;10 Suppl:S10-7.

6. Hornykiewicz O Fau - Kish SJ, Kish SJ. Neurochemical basis of dementia in Parkinson’s disease. (0317-1671 (Print)).

7. Son HG, Altintas O, Kim EJE, Kwon S, Lee SV. Age-dependent changes and biomarkers of aging in Caenorhabditis elegans. Aging Cell. 2019;18(2):e12853.

8. David DC, Ollikainen N, Trinidad JC, Cary MP, Burlingame AL, Kenyon C. Widespread protein aggregation as an inherent part of aging in C. elegans. PLoS Biol. 2010;8(8):e1000450.

9. Glenn CF, Chow DK, David L, Cooke CA, Gami MS, Iser WB, et al. Behavioral deficits during early stages of aging in Caenorhabditis elegans result from locomotory deficits possibly linked to muscle frailty. J Gerontol A Biol Sci Med Sci. 2004;59(12):1251–60.

10. Rennie M, Lin G, Scarlata S. Multiple functions of phospholipase Cβ1 at a glance. Journal of Cell Science. 2022;135(18).

11. Exton JH. Cell signalling through guanine-nucleotide-binding regulatory proteins (G proteins) and phospholipases. [Review] [110 refs]. European Journal of Biochemistry. 1997;243(1-2):10-20.

12. Gilman AG. G proteins: transducers of receptor-generated signals. Annu Rev Biochem. 1987;56:615–49.

13. Hepler JR, Gilman AG. G-proteins. Trends Biochem Sciences. 1992;17:383–7.

14. Pearce KM, Bell M, Linthicum WH, Wen Q, Srinivasan J, Rangamani P, et al. Gαq-mediated calcium dynamics and membrane tension modulate neurite plasticity. Molecular Biology of the Cell. 2020;31(7):683–94.

15. Garwain O, Pearce KM, Jackson L, Carley S, Rosati B, Scarlata S. Stimulation of the Gαq/phospholipase Cβ1 signaling pathway returns differentiated cells to a stem-like state. The FASEB Journal. 2020;34(9):12663–76.

16. Philip F, Guo Y, Aisiku O, Scarlata S. Phospholipase Cβ1 is linked to RNA interference of specific genes through translin-associated factor X. The FASEB Journal. 2012;26(12):4903–13.

17. Philip F, Sahu S, Golebiewska U, Scarlata S. RNA-induced silencing attenuates G protein-mediated calcium signals. The FASEB Journal. 2016;30(5):1958–67.

18. Qifti A, Jackson, L., Singla A, Garwain O, Scarlata S. Stimulation of phospholipase Cbeta1 by Galphaq promotes the assembly of stress granule proteins. Science Signaling. 2021;14(705):eaav1012.

19. Sahu S, Williams L, Perez A, Philip F, Caso G, Zurawsky W, et al. Regulation of the activity of the promoter of RNA-induced silencing, C3PO. Protein Science. 2017;26(9):1807-18.

20. Aisiku OR, Runnels LW, Scarlata S. Identification of a Novel Binding Partner of Phospholipase Cβ1: Translin-Associated Factor X. PLOS ONE. 2010;5(11):e15001.

21. Anderson P, Kedersha N. RNA granules. J Cell Biol. 2006;172(6):803–8.

22. Buchan JR, Parker R. Eukaryotic stress granules: the ins and outs of translation. Mol Cell. 2009;36(6):932–41.

23. Mahboubi H, Stochaj U. Cytoplasmic stress granules: Dynamic modulators of cell signaling and disease. Biochimica et Biophysica Acta (BBA) - Molecular Basis of Disease. 2017;1863(4):884–95.

24. Wheeler JR, Matheny T, Jain S, Abrisch R, Parker R. Distinct stages in stress granule assembly and disassembly. eLife. 2016;5:e18413.

25. Cao X, Jin X, Liu B. The involvement of stress granules in aging and aging-associated diseases. Aging Cell. 2020;19(4):e13136.

26. Jackson L, Rennie M, Poussaint A, Scarlata S. Activation of Gαq sequesters specific transcripts into Ago2 particles. Scientific Reports. 2022;12(1):8758.

27. Ebert RH, 2nd, Shammas Ma Fau - Sohal BH, Sohal Bh Fau - Sohal RS, Sohal Rs Fau - Egilmez NK, Egilmez Nk Fau - Ruggles S, Ruggles S Fau - Shmookler Reis RJ, et al. Defining genes that govern longevity in Caenorhabditis elegans. (0192-253X (Print)).

28. Houthoofd K, Braeckman BP, Lenaerts I, Brys K, De Vreese A, Van Eygen S, et al. Ageing is reversed, and metabolism is reset to young levels in recovering dauer larvae of C. elegans. Experimental Gerontology. 2002;37(8):1015–21.

29. Yang P, Mathieu C, Kolaitis RM, Zhang P, Messing J, Yurtsever U, et al. G3BP1 Is a Tunable Switch that Triggers Phase Separation to Assemble Stress Granules. Cell. 2020;181(2):325–45.e28.

30. Digman MA, Dalal R, Horwitz AF, Gratton E. Mapping the Number of Molecules and Brightness in the Laser Scanning Microscope. Biophys J. 2008;97:2320–32.

31. Namkoong S, Ho A, Woo YM, Kwak H, Lee JH. Systematic Characterization of Stress-Induced RNA Granulation. Mol Cell. 2018;70(1):175–87 e8.

32. Dupuy D, Jovic K, Sterken MG, Grilli J, Bevers RPJ, Rodriguez M, et al. Temporal dynamics of gene expression in heat-stressed Caenorhabditis elegans. Plos One. 2017;12(12).

33. Baronti L, Guzzetti I, Ebrahimi P, Friebe Sandoz S, Steiner E, Schlagnitweit J, et al. Base-pair conformational switch modulates miR-34a targeting of Sirt1 mRNA. Nature. 2020;583(7814):139-44.

34. Broderick JA, Salomon WE, Ryder SP, Aronin N, Zamore PD. Argonaute protein identity and pairing geometry determine cooperativity in mammalian RNA silencing. RNA (New York, NY). 2011;17(10):1858–69.

35. Braeckman BP, Houthoofd K Fau - Vanfleteren JR, Vanfleteren JR. Patterns of metabolic activity during aging of the wild type and longevity mutants of Caenorhabditis elegans. (2152-4041 (Print)).

36. Floyd RA, West M, Hensley K. Oxidative biochemical markers; clues to understanding aging in long-lived species. Experimental Gerontology. 2001;36(4):619–40.

37. Chance B, Schoener B, Oshino R, Itshak F, Nakase Y. Oxidation-reduction ratio studies of mitochondria in freeze-trapped samples. NADH and flavoprotein fluorescence signals. J Biol Chem. 1979;254(11):4764–71.

38. Stringari C, Cinquin A, Cinquin O, Digman MA, Donovan PJ, Gratton E. Phasor approach to fluorescence lifetime microscopy distinguishes different metabolic states of germ cells in a live tissue. Proceedings of the National Academy of Sciences of the United States of America. 2011;108(33):13582–7.

39. Zhen M, Samuel AD. C. elegans locomotion: small circuits, complex functions. Current opinion in neurobiology. 2015;33:117–26.

40. Liu J, Zhang B, Lei H, Feng Z, Liu J, Hsu A-L, et al. Functional Aging in the Nervous System Contributes to Age-Dependent Motor Activity Decline in C. elegans. Cell Metabolism. 2013;18(3):392–402.

41. Brundage L, Avery L Fau - Katz A, Katz A Fau - Kim UJ, Kim Uj Fau - Mendel JE, Mendel Je Fau - Sternberg PW, Sternberg Pw Fau - Simon MI, et al. Mutations in a C. elegans Gqalpha gene disrupt movement, egg laying, and viability. (0896-6273 (Print)).

42. Alcino J. Silva, Jeffrey H. Kogan, Paul W. Frankland a, Kida S. CREB AND MEMORY. Annual Review of Neuroscience. 1998;21(1):127–48.

43. Jospin M, Qi YB, Stawicki TM, Boulin T, Schuske KR, Horvitz HR, et al. A neuronal acetylcholine receptor regulates the balance of muscle excitation and inhibition in Caenorhabditis elegans. PLoS Biol. 2009;7(12):e1000265.

44. Stevenson ME, Bieri G, Kaletsky R, St Ange J, Remesal L, Pratt KJB, et al. Neuronal activation of G(αq) EGL-30/GNAQ late in life rejuvenates cognition across species. Cell Rep. 2023;42(9):113151.

45. Bastiani CA, Gharib S Fau - Simon MI, Simon Mi Fau - Sternberg PW, Sternberg PW. Caenorhabditis elegans Galphaq regulates egg-laying behavior via a PLCbeta-independent and serotonin-dependent signaling pathway and likely functions both in the nervous system and in muscle. (0016-6731 (Print)).

46. Kuo C-T, You G-T, Jian Y-J, Chen T-S, Siao Y-C, Hsu A-L, et al. AMPK-mediated formation of stress granules is required for dietary restriction-induced longevity in Caenorhabditis elegans. Aging Cell. 2020;19(6):e13157.

47. Chen L, Liu B. Relationships between Stress Granules, Oxidative Stress, and Neurodegenerative Diseases. Oxidative Medicine and Cellular Longevity. 2017;2017:10.

48. Saez I, Vilchez D. The Mechanistic Links Between Proteasome Activity, Aging and Age-related Diseases. Curr Genomics. 2014;15(1):38–51.

49. Jackson L, Yerramilli VS, Scarlata S. Live Cell Fluorescence Imaging Shows Neurotransmitter Activation Promotes Aggregation of the Intracellular Domain of Amyloid Precursor Protein. J Membr Biol. 2022;255(4-5):613–22.

50. Ramaswami M, Taylor JP, Parker R. Altered Ribostasis: RNA-Protein Granules in Degenerative Disorders. Cell. 2013;154(4):727–36.

51. Pan C-L, Peng C-Y, Chen C-H, McIntire S. Genetic analysis of age-dependent defects of the Caenorhabditis elegans touch receptor neurons. Proceedings of the National Academy of Sciences. 2011;108(22):9274–9.

52. Brenner S. The genetics of Caenorhabditis elegans. (0016-6731 (Print)).

53. Palikaras K, SenGupta T, Nilsen H, Tavernarakis N. Assessment of dopaminergic neuron degeneration in a C. elegans model of Parkinson’s disease. STAR Protocols. 2022;3(2):101264.

54. Ash PE, Zhang Yj Fau - Roberts CM, Roberts Cm Fau - Saldi T, Saldi T Fau - Hutter H, Hutter H Fau - Buratti E, Buratti E Fau - Petrucelli L, et al. Neurotoxic effects of TDP-43 overexpression in C. elegans. (1460-2083 (Electronic)).

55. Bhaskaran S, Butler Ja Fau - Becerra S, Becerra S Fau - Fassio V, Fassio V Fau - Girotti M, Girotti M Fau - Rea SL, Rea SL. Breaking Caenorhabditis elegans the easy way using the Balch homogenizer: an old tool for a new application. (1096-0309 (Electronic)).

